# UniWave-2: A Hybrid Model for Nucleic Acid Waveform Feature Extraction Enhanced by Fourier and Wavelet Transforms

**DOI:** 10.64898/2026.09.21.753349

**Authors:** Hao Zhang, Yujun Qi, Yulan Feng, Lili Wang

## Abstract

**Motivatio:** Traditional methods primarily rely on statistical features such as k-mers and GC content, making it difficult to capture complex internal relationships within sequences. Deep learning models typically rely on discrete encodings, leading to issues such as information sparsity, dimensional redundancy, and disruption of sequence continuity. Our previous UniWave-1 framework transforms sequences into waveform signals; however, it lacks comprehensive physicochemical integration and computational efficiency.

**Results:** We propose the UniWave-2 feature extraction framework. UniWave-2 incorporates three key advancements: (i) integration of hydrophobicity into the encoding scheme; (ii) enhancement of waveform resolution through Fourier transform, coupled with wavelet transform to precisely capture both local details and global periodic patterns; and (iii) development of a lightweight multi-scale GRU model that facilitates cross-dimensional feature interactions, captures bidirectional temporal dependencies, and integrates long- and short-range patterns with spatial positional awareness, thereby yielding rich feature representations. UniWave-2 achieved competitive performance across multiple genomic tasks, while ISM analysis demonstrated that waveform encoding enables interpretable attribution without relying on gradient information. These results establish UniWave-2 as a lightweight and interpretable framework for nucleic acid sequence analysis.

**Availability:** https://github.com/qcy7226-create/Uniwave.2

## 1 Introduction

The deep integration of artificial intelligence and biological sciences continues to advance, and efficient and rational feature extraction from biological data has become a central issue in the field of bioinformatics (Mullowney, et al., 2023; Vatansever, et al., 2021; Weerarathna, et al., 2023). When dealing with large-scale nucleic acid sequences that contain complex noise and multi-scale structures, how to extract features that are both biologically meaningful and conducive to model learning directly impacts the accuracy and reliability of sequence analysis algorithms. In key tasks such as promoter recognition, epigenetic modification site prediction, and viral mutation classification, traditional methods primarily rely on statistical features such as k-mers (Du, et al., 2020; Zhang, et al., 2024) and GC content, making it difficult to capture the complex internal relationships within sequences. Although current deep learning approaches have substantially improved predictive performance, their encoding and modeling strategies remain subject to inherent limitations. Conventional architectures typically rely on discrete representations such as one-hot encoding (Zhao, et al., 2022), which lack intrinsic physicochemical information of nucleotides and are incompatible with downstream signal processing modules due to their discrete nature. Pretrained models based on Transformers (Nguyen, et al., 2024; Wang, et al., 2024), while exhibiting strong performance, often come with high computational costs, making them expensive to model long sequences and limiting their interpretability (Alsaleh, et al., 2023). Furthermore, advanced models capable of handling ultra-long sequences typically rely on large-scale pretraining, making them difficult to efficiently apply in standard experimental environments. Therefore, despite continuous advances in deep learning methodologies, two fundamental challenges in biological sequence modeling remain unresolved: (i) designing encoding strategies that capture the intrinsic physicochemical properties of nucleotides, and (ii) reducing computational complexity while preserving model representation capacity. Addressing these challenges requires the development of feature extraction approaches that integrate biological relevance, continuous dense representations, and computational efficiency, which remains a critical direction for improving both predictive performance and interpretability.

From an information-structural perspective, nucleic acid sequences can be regarded as discrete transitions among four nucleotides within a four-dimensional representation space (Supplementary Fig. 1, UniWave-1 mapping scheme), exhibiting an intrinsic oscillatory nature in their information organization. Based on this perspective, we previously developed UniWave-1, a nucleic acid feature extraction approach that transforms sequential information into waveform signals (Zhang, et al., 2026). Notably, many biological functions exhibit periodic characteristics, and such periodic patterns are frequently embedded within genomic sequences. Mathematically, periodicity corresponds to oscillatory behavior; therefore, the periodic properties underlying biological functions may be associated with low-frequency components within nucleic acid-derived waveforms. Preliminary numerical mapping experiments with UniWave-1 demonstrated that when nucleotide encoding incorporates intrinsic physicochemical properties, the model can more effectively capture associations between sequence patterns and functional attributes (Supplementary Table 1). Furthermore, waveform-based encoding provides a novel analytical perspective for uncovering functionally relevant patterns embedded within biological sequences.

By abstracting genomic information into waveform signals, UniWave-1 enables efficient sequence feature extraction and achieves an approximately tenfold acceleration compared with Transformer-based models when processing 1,000 bp sequences. Indeed, interpreting nucleic acid sequences as signals and analyzing them using frequency-domain approaches has a long-standing history in computational biology. Voss et al. first revealed long-range fractal correlations and 1/f^β noise in DNA nucleotide sequences through spectral density analysis (Voss, 1992), demonstrating that genomic sequences exhibit measurable spectral properties. Subsequently, Anastassiou et al. systematically applied frequency-domain analysis to the identification of protein-coding regions (Anastassiou, 2000), establishing a foundation for the broader application of signal processing techniques in genomics. Building upon these advances, a variety of signal processing techniques, including Fourier transforms and wavelet transforms, have been extensively employed for exon prediction, promoter identification, and the analysis of periodic patterns in DNA sequences. For example, the WaveDNA method proposed by Ruggeri et al. employs wavelet transforms for DNA sequence analysis. In addition, numerical mapping based on electron–ion interaction potential (EIIP) has also been widely utilized for signal processing analysis of DNA sequences (Nair and Sreenadhan, 2006). Collectively, these studies established a methodological framework that transforms discrete biological sequences into numerical signals and subsequently analyzes them using signal processing techniques. In contrast to these previous approaches, UniWave is distinguished by directly converting sequences into continuous waveform signals rather than discrete numerical vectors, thereby enabling subsequent deep learning modeling within a unified waveform-based framework. However, UniWave-1 still has three major limitations. First, its numerical mapping does not fully incorporate the intrinsic physico-chemical properties of nucleotides. Second, the window-based interpolation strategy may introduce irrelevant variables and incur additional computational costs. Third, the limited compatibility between the model architecture and the encoding scheme restricts effective feature utilization. In this context, we propose UniWave-2, a lightweight hybrid model for nucleic acid waveform feature extraction enhanced by Fourier interpolation and wavelet transformation. This framework integrates nucleotide physicochemical properties with waveform-based signal processing techniques, further improving model interpretability while maintaining high computational efficiency and predictive accuracy.

The principal contributions of this study can be summarized in five aspects. First, we found that nucleotide hydrophobicity is closely associated with DNA secondary structure and protein-binding affinity. To address this, we developed a physicochemical property-based encoding framework using nucleotide hydrophobicity, in which the four nucleotides are mapped into continuous numerical values. This representation not only provides a degree of biological interpretability but also ensures compatibility with signal processing frameworks, thereby addressing the limitation of discrete encoding schemes, such as DNABERT (Ji, et al., 2021) and ChromBERT (Lee, et al., 2024), which lack explicit physicochemical information. Second, we design a processing pipeline termed “Fourier-interpolation signal densification followed by wavelet convolution–based compression and denoising.” Specifically, Fourier zero-padding interpolation (Mottola, et al., 2021) is applied to increase signal density, followed by sym6 wavelet decomposition and adaptive signal enhancement, thereby strengthening key functional features while reducing sequence redundancy and maintaining a balance between information retention and computational efficiency. Third, we design a task-adaptive deep learning architecture, MultiScaleGRU. This design effectively integrates high-dimensional representations, multi-scale spatial features, bidirectional temporal dependencies, and positional awareness, providing rich feature representations for downstream tasks. Fourth, the waveform encoding framework provides a distinctive perspective for deep learning model research from the standpoint of physical wave representations, offering a valuable approach for network visualization and interpretability analysis. Finally, we establish an evaluation framework comprising eight datasets spanning four task categories—viral variant classification, promoter identification, m6A site prediction, and subspecies typing. The experimental results indicate that UniWave-2 exhibits strong generalizability and robustness across diverse nucleic acid information processing tasks, highlighting its broad application potential in bioinformatics and downstream artificial intelligence–driven analyses.

## 2 Methods

The overall UniWave framework is divided into two components: a continuous waveform encoding (Fig. 1) and the end-to-end architecture of MultiScaleGRU (Fig. 2). Data transfer between these components is co-ordinated through H5 files that enable efficient data storage and retrieval.

**Fig. 1.**
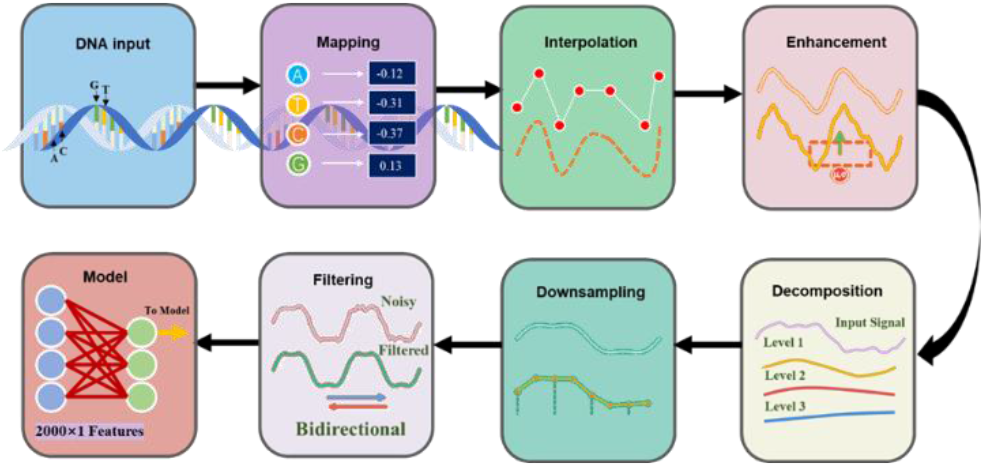
Continuous waveform encoding. In this pipeline, DNA bases are first mapped to hydrophobicity-based numerical signals. Fourier interpolation is then applied to reconstruct the temporal sequence, followed by adaptive signal enhancement to improve feature discriminability. Subsequently, wavelet decomposition and downsampling are performed to achieve multi-scale signal compression, after which bidirectional mean filtering is applied for noise reduction. The final output is a N × 1 feature representation used as the input to the model.

**Fig. 2.**
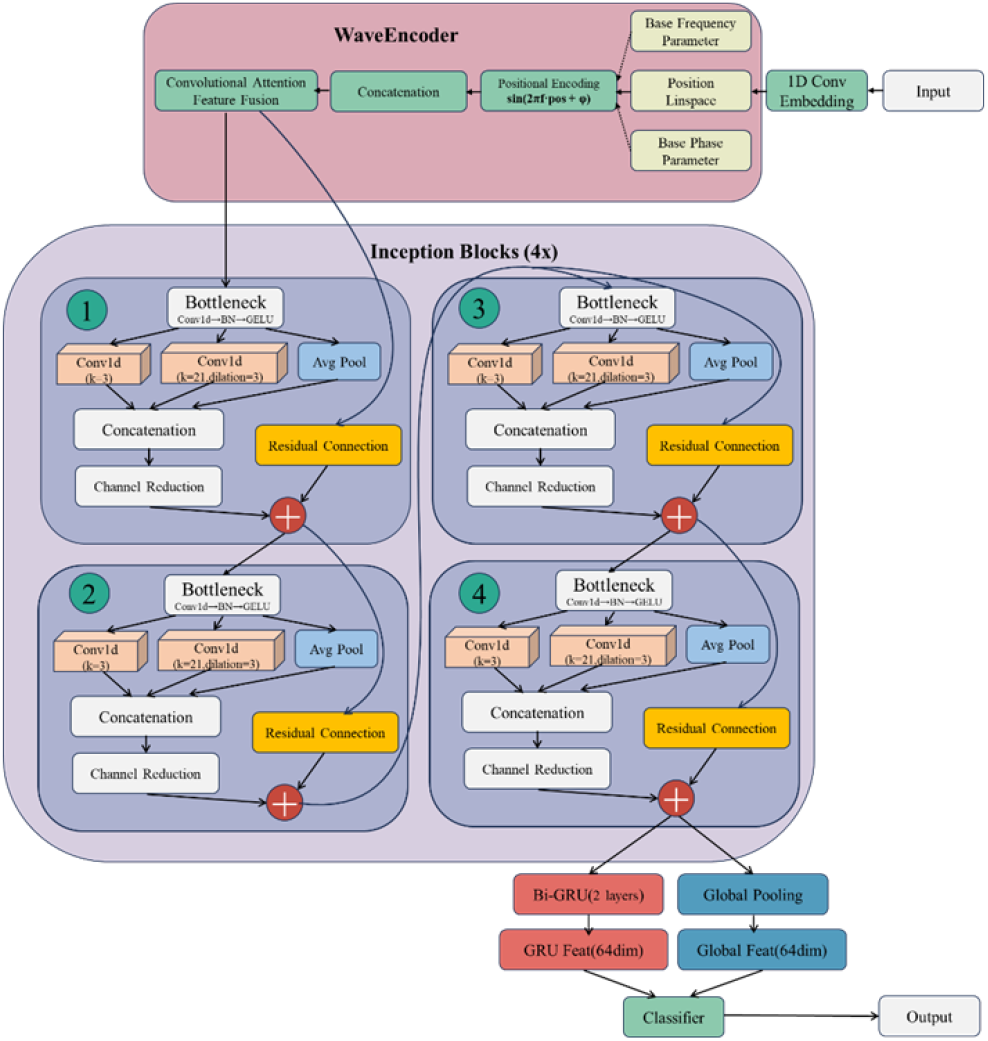
End-to-end architecture of MultiScaleGRU. The model takes DNA sequences as input. After 1D convolutional embedding, the representation is first enhanced by the WaveEncoder, which integrates learnable base-frequency/phase sinusoidal positional encoding with convolutional attention to strengthen temporal positional awareness. The features are then passed through four stacked Inception Blocks (incorporating bottleneck compression and residual connections) to achieve multi-scale feature extraction. Finally, a bidirectional GRU captures long-range temporal dependencies; the resulting representations are fused with global pooling features and fed into a classifier to produce the final predictions.

### 2.1 Construction of a novel continuous waveform encoding

Conventional discrete encoding strategies mainly include one-hot encoding and k-mer embedding. The former represents A, T, C, and G as four-dimensional one-hot vectors (e.g., A→[1,0,0,0]), completely disregarding the physicochemical differences among nucleotides. The latter relies on sliding-window-based k-mer frequency statistics, resulting in the loss of fine-grained information at the single-nucleotide level. Pre-trained models represented by DNABERT-2 tokenize sequences into 6–10 bp segments and generate 768-dimensional abstract embeddings. Although these models can capture long-range dependencies, their tokenization strategies may disrupt the integrity of regulatory elements, such as short motifs including TATA-box sequences. In contrast, our waveform encoding strategy does not require sequence segmentation but instead directly transforms the entire DNA sequence into a continuous waveform representation, inherently preserving sequence integrity and continuity. Although other physicochemical properties, such as hydrogen bonding and electrostatic interactions, also contribute substantially to nucleic acid structure and function, the particular value of hydrophobicity lies in its ability to characterize physicochemical differences arising from nucleotide composition and its association with base-stacking stability and nucleic acid–molecule interactions (Boldina, et al., 2009). Therefore, hydrophobicity provides a biophysically meaningful continuous representation that links sequence information with structural features and molecular interaction states. Based on this rationale, we selected nucleotide hydrophobicity as a key physicochemical property underlying DNA structure and function, providing a natural biological anchor for sequence encoding.

To establish a nucleic acid hydrophobicity scale with explicit physicochemical significance, we employed the logP values (octanol–water partition coefficients) of the four nucleotide bases calculated using the Ghose– Crippen atomic contribution method (Ghose and Crippen, 1986; Guckian, et al., 2000). LogP is a classical thermodynamic descriptor of molecular hydrophobicity that reflects the partitioning preference of molecules between aqueous and organic phases. This method has been extensively validated and widely applied in medicinal chemistry and computational biology (Wildman and Crippen, 1999).Consequently, a hydrophobicity-derived waveform representation emerges, establishing a direct link between encoded features and molecular function. The core advantages of this design are threefold: (1) preservation of hydrophobicity differences, allowing direct reflection of base-pair stability and protein-binding potential; (2) Reducing the feature dimensions to a single dimension results in approximately a 60% reduction in computational cost compared to DNABERT-2; and (3) the continuous numerical representation can be directly converted into time-domain signals without additional format transformation.

Through numerical mapping, we obtain a hydrophobicity-based time-domain signal, which can then be transformed into a waveform representation.

According to the Shannon Sampling Theorem, a band-limited signal (bandwidth ≤ B Hz) can be uniquely reconstructed from its discrete samples, with the ideal reconstruction achieved using a sinc interpolation kernel. Based on this principle, the discrete sequence *x*[*n*] is first transformed from the time domain to the frequency domain via the discrete Fourier transform (DFT). In the frequency domain, zero-padding is applied to the high-frequency components of the spectrum to extend its length and increase the sampling resolution in the frequency domain. Subsequently, an inverse discrete Fourier transform (IDFT) is performed on the zero-padded spectrum to obtain a new time-domain signal *x*^′^[*n*]. The resulting signal *x*^′^[*n*] effectively represents a smooth and continuous approximation of the original signal obtained through sinc interpolation. In other words, zero-padding in the frequency domain serves as an efficient implementation of the ideal interpolation process (See Supplementary Figure 2). This strategy also addresses the issue in UniWave-1 where window-function interpolation introduces irrelevant variables, while substantially improving computational efficiency.

Through this process, the waveform is rendered continuous, resulting in a one-dimensional continuous signal derived from hydrophobicity values. However, simple waveform transformation alone does not sufficiently enhance informative features within the sequence, limiting the model’s ability to effectively learn biologically relevant patterns. In practice, feature extraction would still largely depend on neural networks, while Fourier interpolation increases sequence length, which is inconsistent with our goal of maintaining a lightweight computational framework. Therefore, a method is required that can simultaneously perform downsampling while enhancing the extraction of critical sequence features. Conventional downsampling techniques often lead to substantial information loss from the original sequence, which is detrimental for analyzing both local patterns and long-range dependencies. To address this limitation, we adopt the wavelet transform, a signal-processing technique closely related to the Fourier transform. Wavelet methods (Michau, et al., 2022) have been widely applied across multiple domains, and their integration with deep learning frameworks has recently gained increasing attention. Leveraging the inherent signal decomposition capability of wavelet transforms, we develop the WCJC method to perform multi-scale decomposition of oversampled signals, separating global trends and local features. The signal is then reconstructed by combining one global component with multiple local components, followed by threefold downsampling of the reconstructed signal (detailed procedures are provided in the Supplementary Materials). This strategy effectively preserves key features while reducing sequence length. Moreover, the separation and reconstruction of key signal components facilitate secondary extraction of embedded biochemical properties and structural characteristics within biological sequences.

To further enhance highly variable regions—such as those containing functional motifs—while moderately amplifying stable regions, we design an adaptive waveform enhancement and local feature amplification scheme following the interpolation stage. In highly fluctuating regions, local normalization is first applied, after which the signal is multiplied by a gain factor of 0.2 and superimposed onto the original signal to emphasize feature differences. In relatively stable regions, the original signal is directly scaled by a factor of 1.1 to achieve moderate amplification. This strategy suppresses noise while emphasizing biologically relevant signals, thereby improving the intrinsic robustness of the encoded data. Finally, a lightweight filtering step using a finite impulse response (FIR) filter (Rui, et al., 2024) is applied to the WCJC-processed waveform to suppress potential high-frequency noise remaining after downsampling. A weighted fusion of the smoothed waveform and the original signal is then performed to preserve local fine-grained features, thereby preventing excessive smoothing and the loss of critical information. Through these steps, a complete waveform-based feature extraction pipeline is established, providing high-quality feature inputs for subsequent model training. The above processing steps were performed sequentially as follows: Fourier interpolation, adaptive enhancement, wavelet decomposition and downsampling, followed by FIR filtering. Another key advantage of this waveform encoding pipeline is its inherent interpretability. Because waveform signals preserve the spatial continuity of the original sequence, functional importance at individual positions can be directly assessed through sequence-level perturbation analysis on the original sequences, without relying on gradient-based attribution methods.

### 2.2 The MultiScaleGRU hybrid model

To address key challenges in genomic signal analysis—including the co-existence of multi-scale functional motifs, pronounced bidirectional temporal dependencies, and the strong association between gene function and positional context—we design a deep learning model termed MultiScaleGRU. MultiScaleGRU first employs a convolutional embedding layer to project the one-dimensional signal into a higher-dimensional feature space, providing sufficient representational capacity for subsequent feature extraction while maintaining signal continuity. Since genomic signal functionality is closely related to its positional context, traditional fixed-position encoding schemes cannot dynamically adjust positional weights to different functional regions. To enhance positional awareness, we propose the attention-guided position encoding module, WaveEncoder, which dynamically integrates positional information with waveform features. Subsequently, we introduce the improved InceptionTime module, incorporating dilated convolutions to expand the receptive field and improve multi-scale feature extraction efficiency, enabling simultaneous capture of key spatial features from local short sequence motifs and global long-range regulatory regions. Finally, a bidirectional GRU (Cho, et al., 2014) layer models the inherent 5′→3′ and 3′→5′ bidirectional dependencies of double-stranded DNA, and the fused features are fed into a classifier to perform task-specific predictions.

MultiScaleGRU contains only about 267,000 parameters, less than 1% of the size of mainstream genomic foundation models, and can efficiently process 2000 bp sequences on consumer-grade GPUs with less than 8 GB of memory, achieving a good balance between prediction accuracy and computational efficiency. The detailed structure and module design of the model can be found in the supplementary materials.

### 2.3 Evaluation strategy

To meet the standard requirements for deep learning evaluation, we stratified the data by category and divided each task into a training set (70%), a validation set (20%), and an independent test set (10%). The training set is used for model parameter learning, the validation set is used for early stopping and hyperparameter tuning, and the independent test set is used for a final evaluation only after the model training is complete, without being involved in any training or validation decisions during the process. All final experimental results were reported on independent test sets, including balanced accuracy (BA), macro-averaged F1 score (Macro-F1), AUC-ROC, and their corresponding 95% confidence intervals estimated using 1,000 bootstrap iterations, rather than relying on averaged errors from cross-validation. For experiments involving comparisons with published benchmark models on public datasets, we prioritized the use of the same evaluation metrics reported in the original benchmark studies to ensure fair comparisons. The specific metrics used for each task are indicated in the corresponding tables.

## 3 Results

We systematically evaluate UniWave-2 using established benchmark classification tasks, which include multiple publicly available datasets and experimental results from mainstream models, in order to comprehensively investigate its versatility and computational advantages across diverse tasks involving multi-species genomes.

### 3.1 Dataset

All datasets used in this study are constructed from publicly available benchmark resources, following the principles of reproducibility and standardized evaluation, and are designed to support different task-specific assessment requirements. The multi-species promoter recognition dataset is derived from Deepromclass (Kari, et al., 2023), covering five organisms—Drosophila, yeast, mouse, C. elegans, and human—with sequence length gradients of 80 bp, 150 bp, and 300 bp. This dataset is used to evaluate the model’s cross-species generalization ability and adaptability to different sequence lengths. The rice m6A site identification dataset (Wang, et al., 2024), a widely recognized benchmark in the field, contains experimentally validated m6A-modified positive sites and non-modified negative sites, enabling evaluation of the model’s ability to capture conserved motifs associated with epigenetic modifications. The SARS-CoV-2 nine-variant classification dataset, derived from the DNABERT-2 benchmark, contains 1000 bp sequences from nine variants, including Alpha and Beta. It is designed to assess the model’s capability for long-sequence modeling and mutation pattern recognition. Only a single challenging test set is used, which includes nucleotide mutations, base pair fragment insertions/deletions, and Gaussian noise, to evaluate the model’s robustness under complex perturbations. Specific perturbation metrics are provided in the supplementary materials.

### 3.2 UniWave-2 efficiently identifies promoter regions across multiple species

Promoter recognition is a fundamental task for elucidating gene transcriptional regulatory mechanisms, and its performance directly influences the accuracy of downstream molecular biology studies. To systematically validate the effectiveness of the proposed model, we conducted a comprehensive evaluation of promoter data from five eukaryotic species using the publicly available Deepromclass dataset framework. The model was further compared with three representative baseline models provided in the dataset—Inception, ResNet, and CNN+BiLSTM— as shown in Table 1.

**Table 1.** Performance comparison of promoter recognition across five eukaryotic species using different sequence lengths.

| Species | Model | Sequence Length (bp) |  |  |
| --- | --- | --- | --- | --- |
|  |  | 80 | 150 | 300 |
| Drosophila | Inception | / | 0.792 | / |
|  | Resnet | / | 0.894 | / |
|  | CNN+BiLSTM | 0.912 | 0.918 | 0.927 |
|  | UniWave-2 | **0.936** | **0.940** | **0.958** |
| Yeast | Inception | / | 0.811 | / |
|  | Resnet | / | 0.700 | / |
|  | CNN+BiLSTM | **0.893** | 0.907 | 0.909 |
|  | UniWave-2 | 0.826 | **0.909** | **0.942** |
| Mouse | Inception | / | 0.784 | / |
|  | Resnet | / | 0.850 | / |
|  | CNN+BiLSTM | 0.861 | 0.865 | 0.897 |
|  | UniWave-2 | **0.869** | **0.907** | **0.927** |
| *C. elegans* | Inception | / | / | / |
|  | Resnet | / | 0.884 | / |
|  | CNN+BiLSTM | **0.898** | 0.936 | 0.924 |
|  | UniWave-2 | 0.876 | **0.939** | **0.942** |
| Human | Inception | / | 0.728 | / |
|  | Resnet | / | 0.833 | / |
|  | CNN+BiLSTM | 0.831 | 0.840 | 0.855 |
|  | UniWave-2 | **0.851** | **0.883** | **0.891** |

The evaluation metric used is the AUC–ROC score, which is well suited for binary classification tasks. The results show that, across nearly all species and sequence length settings, UniWave-2 consistently outperforms all baseline models, with the only exception occurring at the 80 bp sequence length for yeast and C. elegans, where CNN+BiLSTM achieves slightly higher scores.

In the sequence length dependency analysis, the performance of Uni-Wave-2 shows a consistent improvement as the sequence length increases. Under the 300-base-pair sequence setting, compared with the best-performing CNN+BiLSTM model at the same sequence length, the recognition accuracy for the five species increased by 0.031, 0.033, 0.030, 0.018, and 0.036, respectively.

Cross-species performance evaluation indicates that UniWave-2 demonstrates robust performance across eukaryotic organisms with varying evolutionary distances. For model organisms such as C. elegans and Drosophila, the recognition accuracy generally exceeds 0.93. For organisms with complex genomes, such as humans, the accuracy still reaches 0.891, representing improvements of 0.058 and 0.036 over ResNet (0.833) and CNN+BiLSTM (0.855), respectively. These findings suggest that UniWave-2 can effectively capture both conserved promoter features across species and species-specific sequence patterns.

### 3.3 Identification of m6A sites in rice

N6-methyladenosine (m6A) is one of the most abundant methylation modifications in eukaryotic RNA and plays a critical regulatory role in rice growth, development, and stress response. Accurate identification of m6A sites in rice is therefore an essential prerequisite for elucidating its epigenetic regulatory mechanisms. To evaluate the performance of UniWave-2 on this task, we conducted experiments based on a publicly available benchmarking framework for rice m6A site identification. The model was compared with several state-of-the-art tools—iDNA6mA-Rice, SNN-Rice6mA, SpineNet-6mA, and i6mA-CNN—as well as UniWave-1, previously developed by our team. Accuracy (ACC), area under the receiver operating characteristic curve (AUC), and confidence intervals were used as the primary evaluation metrics (Table 2).

**Table 2.** Comparison with state-of-the-art methods for rice m6A site identification.

| Tools | BA | AUC-ROC | Accuracy (95% CI) |
| --- | --- | --- | --- |
| iDNA6mA-Rice | 91.7% | 0.96 | / |
| SNNRice6mA | 92.0% | 0.97 | / |
| Uniwave-1 | 92.16% | 0.97 | 95% CI: 91.94%–92.36% |
| SpineNet-6mA | 94.31% | 0.98 | / |
| i6mA-CNN | 93.97% | 0.98 | / |
| **Uniwave-2** | **94.68%** | **0.98** | **95% CI: 94.50%–94.88%** |

The experimental results demonstrate that UniWave-2 achieves outstanding overall performance in the rice m6A site identification task, ranking among the top across all evaluation metrics. In terms of the key accuracy (ACC) metric, UniWave-2 reaches 94.68%, significantly outperforming conventional mainstream tools. Compared with UniWave-1, the previous model in the same series, UniWave-2 improves accuracy from 92.16% to 94.68%, representing a 2.52 percentage point increase, which highlights the effectiveness of the model’s iterative optimization. In terms of the AUC metric, UniWave-2 achieved a high score of 0.98, demonstrating strong discriminative ability between positive and negative samples. In addition, UniWave-2 reports a 95% confidence interval for accuracy (94.50%–94.88%), further validating the stability and reliability of its predictions. This characteristic provides a more reliable reference for subsequent experimental validation.

Overall, UniWave-2 achieves both breakthrough accuracy and strong result stability in the rice m6A site identification task.

### 3.4 Classification task of nine SARS-CoV-2 variants

This section evaluates the performance of UniWave-2 based on the SARS-CoV-2 variant classification task framework introduced in the DNABERT-2 study. The dataset used in this task contains sequences of 1000 bp, serving as a benchmark for assessing the model’s long-sequence modeling capability. It should be noted that all baseline models used for comparison—including the DNABERT series, the Nucleotide Transformer series, and DNABERT-2—are foundation models pretrained on multi-species or human genomic data. In contrast, UniWave-2 is trained using an end-to-end framework without any pretraining. This fundamental difference in training paradigms should be considered when interpreting the results (Table 3).

**Table 3.** Performance comparison with large-scale pretrained foundation models on SARS-CoV-2 variant classification.

| Model | Num. Params | Macro-F1 |
| --- | --- | --- |
| DNABERT (3-mer) | 86M | 62.23 |
| DNABERT (4-mer) | 86M | 59.87 |
| DNABERT (5-mer) | 87M | 63.64 |
| DNABERT (6-mer) | 89M | 55.50 |
| NT-500M-human | 480M | 57.13 |
| NT-500M-1000g | 480M | 52.06 |
| NT-2500M-1000g | 2537M | 66.73 |
| NT-2500M-multi | 2537M | 73.04 |
| DNABERT-2 | 117M | 71.02 |
| DNABERT-2† | 117M | 68.49 |
| **UniWave-2** | **0.33M** | **57.46** |

Note: †Indicates that the model underwent additional masked language model (MLM) pre-training on the training sets of the GUE benchmark.

The experiment uses the F1 score as the primary evaluation metric, while also considering the model parameter size to assess computational efficiency. Experimental results show that the F1 score of UniWave-2 is lower than that of large pretrained models such as NT-2500M-multi (73.04) and DNABERT-2 (71.02). However, it still surpasses several pretrained models, outperforming NT-500M-1000g (52.06) and DNABERT (6-mer) (55.50), while achieving performance comparable to NT-500M-human (57.13). Considering that UniWave-2 does not rely on a pretraining stage for parameter accumulation, and its parameter size is only 1/1454 of NT-500M-human and 1/354 of DNABERT-2, these results highlight its remarkable parameter efficiency. With extremely low model complexity, it achieves classification performance comparable to that of several large pretrained models.

Further analysis indicates that pretrained models, leveraging prior knowledge learned from large-scale genomic datasets, demonstrate superior performance in viral variant classification tasks. However, this advantage comes at the cost of substantial pretraining computational resources and large model storage requirements. In contrast, UniWave-2 directly learns task-specific features through end-to-end training, achieving competitive performance with a lightweight architecture without requiring pretraining. This provides a new approach for rapid SARS-CoV-2 variant detection in resource-constrained environments.

### 3.5 Dengue virus subtype classification

The limited cross-protective immunity among dengue virus (DENV) serotypes increases the risk of severe disease; therefore, rapid and accurate serotype classification is critical for effective outbreak control and clinical management. The dengue virus serotype classification task is relatively straightforward, as the four serotypes exhibit distinct genomic signatures. Therefore, within this experimental framework, this task is not intended to demonstrate conventional classification performance, but rather serves as a benchmark for evaluating model robustness under challenging perturbations. The challenge test set simulates potential sequence variations encountered in real-world scenarios, including nucleotide substitutions, fragment insertions/deletions, and Gaussian noise, to systematically evaluate the robustness of different encoding strategies against perturbations.

The experimental results demonstrate that UniWave-2 exhibits outstanding classification performance and robustness on the challenging test set (Table 4). Compared with One-hot, Embedding, and UniWave-1, all evaluation metrics show substantial improvements, validating the effectiveness of Fourier interpolation as well as the strong compatibility between the MultiScaleGRU hybrid architecture and the waveform encoding scheme.

**Table 4.** Robustness evaluation on a challenge test set for dengue virus serotype classification.

| Encoding Method | Performance on the Challenging Test Set |  |  |  |
| --- | --- | --- | --- | --- |
|  | Macro-F1 | BA | AUC-ROC | Accuracy (95% CI) |
| One-hot | 0.7778 | 77.51% | 0.9563 | 0.7534-0.7981 |
| Embedding | 0.7416 | 72.41% | 0.9805 | 0.7036-0.7458 |
| UniWave-1 | 0.9222 | 92.22% | 0.9917 | 0.9082-0.9358 |
| **UniWave-2** | **0.9950** | **99.50%** | **0.9974** | **0.9912~0.9984** |

### 3.6 Ablation study

To quantify the contribution of each core component of UniWave-2—including the embedding layer, WaveEncoder, GRU, and dilated convolution—and to validate the rationality of the overall architecture design, we conducted an ablation study based on the previously described classification task framework. The performance of each ablation variant is comprehensively compared with that of the complete model (All) to clarify the functional importance of each component (Table 5). The ablation experiments were conducted on the SARS-CoV-2 variant classification task.

**Table 5.** Ablation study of UniWave-2 components.

| Tools | Macro-F1 | BA | AUC-ROC | Accuracy (95% CI) |
| --- | --- | --- | --- | --- |
| All | 0.5801 | 0.5820 | 0.9284 | 0.5727~0.5906 |
| Re embedding | 0.5587 | 0.5610 | 0.9201 | 0.5471~0.5739 |
| Re WaveEncoder | 0.5308 | 0.5302 | 0.9109 | 0.5171~0.5439 |
| Re GRU | 0.5448 | 0.5477 | 0.9212 | 0.5340~0.5611 |
| Re Dilated Convolution | 0.4659 | 0.4673 | 0.8911 | 0.4535~0.4806 |

Note: ‘All’ represents the full UniWave-2 model. ‘Re X’ denotes the model with component X removed (e.g., Re embedding: without embedding layer; Re WaveEncoder: without WaveEncoder; Re GRU: without GRU; Setting the dilation rate to 1, thereby eliminating the receptive field expansion introduced by dilated convolutions).

The experimental results indicate that the full model achieves the best overall performance. All ablation variants exhibit performance degradation to varying degrees. Removing dilation rates resulted in the most pronounced performance degradation, indicating that dilation rates in dilated convolutions substantially contribute to performance improvements under a comparable computational parameter budget. Removing either the WaveEncoder or the GRU also leads to noticeable decreases across all evaluation metrics, confirming their critical roles in temporal positional awareness and in capturing sequential dependencies. The performance decline caused by removing the 1D convolutional embedding layer is relatively moderate, yet still inferior to that of the full model, suggesting that the embedding layer facilitates high-dimensional feature mapping and enhances information from the original features. Overall, the integrated architecture combining embedding, WaveEncoder, GRU, and dilated convolution achieves efficient complementarity across multiple feature dimensions, jointly establishing a comprehensive and robust feature representation framework.

### 3.7 Interpretability Experiments

To investigate the sequence features learned by the waveform encoding model and further elucidate the potential biological basis underlying its predictions, we performed in silico saturation mutagenesis (ISM) to assess model interpretability. Unlike attribution methods that rely on end-to-end gradient propagation, ISM is a model-agnostic perturbation-based approach that systematically introduces nucleotide substitutions within sliding windows across the original DNA sequence and measures the resulting changes in prediction probabilities, thereby quantifying the relative contribution of individual sequence positions to model predictions. We performed ISM analysis on promoter recognition tasks across five species and obtained species-specific positional importance profiles (Fig. 3a–e).

**Fig. 3.**
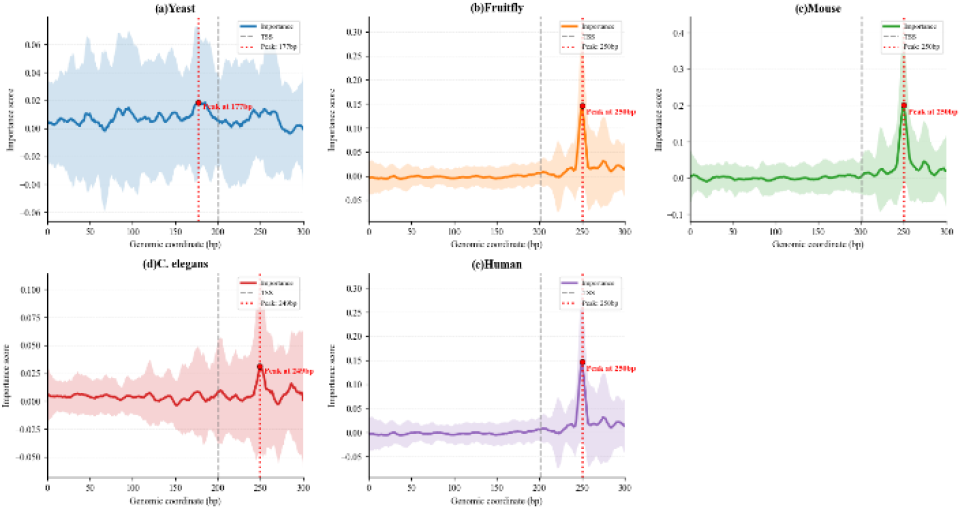
ISM-based positional importance profiles across species. (a–e) ISM importance profiles for the promoters of Saccharomyces cerevisiae, Drosophila melanogaster, Mus musculus, Caenorhabditis elegans, and Homo sapiens, respectively. The gray dashed line indicates the transcription start site (TSS; 201 bp). Red-shaded regions indicate the major importance peak regions. The yeast promoter exhibits a major peak at approximately 170 bp, corresponding to a position approximately 31 bp upstream of the TSS and adjacent to the canonical TATA-box region. The other four species exhibit major peaks at approximately 250 bp, corresponding to a position approximately 49 bp downstream of the TSS and located within the downstream promoter region. Shaded regions indicate ±1 standard deviation (SD).

ISM analysis revealed species-specific positional importance patterns across promoter sequences from different species. As shown in Fig. 3, the ISM importance profile of the yeast promoter exhibited a prominent peak at approximately 170 bp, corresponding to a region approximately 31 bp upstream of the TSS and coinciding with the typical core promoter region containing the TATA box. In contrast, the major importance peaks in the promoters of Drosophila melanogaster, Mus musculus, Caenorhabditis elegans, and Homo sapiens were all located at approximately 250 bp, corresponding to a position approximately 49 bp downstream of the TSS. This peak falls within the downstream core promoter region, indicating that UniWave-2 identified regions associated with downstream-of-TSS sequences that make substantial contributions to its predictions in these species. Because specific core promoter elements were not independently annotated or coordinate-validated in this study, we do not directly attribute this peak to any particular element, such as the DPE. Instead, we interpret it as a species-specific downstream promoter sequence feature identified by the model. Notably, yeast promoters typically rely heavily on core promoter elements such as the TATA box, whereas promoters in many higher eukaryotes can utilize other core promoter elements, including the downstream promoter element (DPE), to mediate transcription initiation (Kutach and Kadonaga, 2000). Without being provided with prior information regarding species identity or the positions of specific promoter elements, UniWave-2 nevertheless produced importance profiles with distinct positional differences across speciesThese findings suggest that the waveform features learned by the model may capture more than nucleotide composition alone and may encode sequence information associated with species-specific organization of promoter regions.

The unique advantage of waveform encoding lies in its ability to map nucleotide substitutions into continuous changes in hydrophobicity values, enabling ISM perturbations to simulate biologically relevant physicochemical alterations. Meanwhile, waveform encoding preserves sequence continuity through Fourier interpolation and wavelet transformation, allowing functional elements (such as the ~7 bp TATA-box motif) to appear as contiguous regions rather than isolated positions in importance profiles. This enables the model to sensitively capture subtle differences in promoter sequences across species. This sequence–wave-form–function mapping provides a new computational perspective for understanding species-specific regulation of promoter activity.

## 4 Discussion

In this study, we present UniWave-2, a lightweight waveform encoding-based model for nucleic acid sequence analysis. By converting discrete DNA sequences into continuous waveform signals through hydrophobicity-based numerical mapping, UniWave-2 achieves competitive predictive performance across diverse species and tasks while maintaining low computational costs. More importantly, its waveform encoding paradigm enables gradient-free ISM perturbation analysis, providing a new technical avenue for improving the interpretability of deep learning models in genomics without requiring large-scale pretraining. Compared with the first-generation UniWave-1 model, UniWave-2 demonstrates substantial improvements in interpretability, predictive accuracy, robustness, and computational efficiency.

Future research directions include the following: (1) The anchors used for waveform derivation will be extended to more fields. (2) Extending the model to more epigenetic regulation tasks, such as m5C methylation site identification and RNA editing site prediction, to further enhance its cross-task generalization capability. (3) Continuously optimizing the model architecture and training strategies, gradually moving toward a lightweight pretraining paradigm to support a broader range of genomic analysis scenarios. (4) Exploring the integration of the model with protein sequence analysis, with a particular focus on DNA–protein interaction prediction.

## References

Alsaleh, M.M., et al. Prediction of disease comorbidity using explainable artificial intelligence and machine learning techniques: A systematic review. Int J Med Inform 2023;175:105088.

Anastassiou, D. Frequency-domain analysis of biomolecular sequences. Bioinformatics 2000;16(12):1073–1081.

Boldina, G., Ivashchenko, A. and Régnier, M. Using Profiles Based on Nucleotide Hydrophobicity to Define Essential Regions for Splicing. International Journal of Biological Sciences 2009;5(1):13–19.

Cho, K., et al. Learning Phrase Representations using RNN Encoder–Decoder for Statistical Machine Translation. In. Doha, Qatar: Association for Computational Linguistics; 2014. p. 1724–1734.

Du, Z., et al. DeepAdd: Protein function prediction from k-mer embedding and additional features. Comput Biol Chem 2020;89:107379.

Ghose, A.K. and Crippen, G.M. Atomic Physicochemical Parameters for Three-Dimensional Structure-Directed Quantitative Structure-Activity Relationships I. Partition Coefficients as a Measure of Hydrophobicity. Journal of Computational Chemistry 1986;7(4):565–577.

Guckian, K.M., et al. Factors Contributing to Aromatic Stacking in Water: Evaluation in the Context of DNA. J Am Chem Soc 2000;122(10):2213–2222.

Ji, Y., et al. DNABERT: pre-trained Bidirectional Encoder Representations from Transformers model for DNA-language in genome. Bioinformatics 2021;37(15):2112–2120.

Kari, H., et al. DeePromClass: Delineator for Eukaryotic Core Promoters Employing Deep Neural Networks. IEEE/ACM Trans Comput Biol Bioinform 2023;20(1):802–807.

Kutach, A.K. and Kadonaga, J.T. The downstream promoter element DPE appears to be as widely used as the TATA box in Drosophila core promoters. Mol Cell Biol 2000;20(13):4754–4764.

Lee, S., et al. ChromBERT: Uncovering Chromatin State Motifs in the Human Genome Using a BERT-based Approach. bioRxiv 2024:2024.2007.2025.605219.

Michau, G., Frusque, G. and Fink, O. Fully learnable deep wavelet transform for unsupervised monitoring of high-frequency time series. Proc Natl Acad Sci U S A 2022;119(8).

Mottola, M., et al. Reproducibility of CT-based radiomic features against image resampling and perturbations for tumour and healthy kidney in renal cancer patients. Sci Rep 2021;11(1):11542.

Mullowney, M.W., et al. Artificial intelligence for natural product drug discovery. Nat Rev Drug Discov 2023;22(11):895–916.

Nair, A.S. and Sreenadhan, S.P. A coding measure scheme employing electron-ion interaction pseudopotential (EIIP). Bioinformation 2006;1(6):197–202.

Nguyen, E., et al. Sequence modeling and design from molecular to genome scale with Evo. Science 2024;386(6723):eado9336.

Rui, L., et al. Signal processing collaborated with deep learning: An interpretable FIRNet for industrial intelligent diagnosis. Mechanical Systems and Signal Processing 2024;212:111314.

Vatansever, S., et al. Artificial intelligence and machine learning-aided drug discovery in central nervous system diseases: State-of-the-arts and future directions. Med Res Rev 2021;41(3):1427–1473.

Voss, R.F. Evolution of long-range fractal correlations and 1/f noise in DNA base sequences. Physical Review Letters 1992;68(25):3805–3808.

Wang, G., et al. Quantitative profiling of m6A at single base resolution across the life cycle of rice and Arabidopsis. Nature Communications 2024;15(1):4881.

Wang, X., et al. A pathology foundation model for cancer diagnosis and prognosis prediction. Nature 2024;634(8035):970–978.

Weerarathna, I.N., Kamble, A.R. and Luharia, A. Artificial Intelligence Applications for Biomedical Cancer Research: A Review. Cureus 2023;15(11):e48307.

Wildman, S.A. and Crippen, G.M. Prediction of Physicochemical Parameters by Atomic Contributions. Journal of Chemical Information and Computer Sciences 1999;39(5):868–873.

Zhang, H., Qi, Y. and Wang, L. UniWave: A Waveform-Based Encoding Framework for Nucleic Acid Feature Extraction. bioRxiv 2026:2026.2001.2012.698567.

Zhang, W., et al. Improving plant miRNA-target prediction with self-supervised k-mer embedding and spectral graph convolutional neural network. PeerJ 2024;12:e17396.

Zhao, J., et al. CNNArginineMe: A CNN structure for training models for predicting arginine methylation sites based on the One-Hot encoding of peptide sequence. Front Genet 2022;13:1036862.

