## Supplementary for "UniWave-2: A Hybrid Model for Nucleic Acid Waveform Feature Extraction Enhanced by Fourier and Wavelet Transforms"

### Supplementary theory:

#### Model Architecture Design:

High-dimensional transformation via a 1D convolutional embedding layer. The raw representation of genomic signals is a continuous numerical sequence (derived from waveform encoding) that lacks the natural "word boundaries" found in natural language. Traditional discrete token embeddings (e.g., k-mer mapping) disrupt the continuity of the signal, whereas directly feeding low-dimensional signals into the model is insufficient to support complex feature extraction. To address this issue, the model incorporates a 1D convolution-based embedding layer as a core component for input adaptation. This layer employs a convolutional kernel size of 3 bp—corresponding to the typical minimum length of functional motifs in genomic sequences, such as the core segments of CpG islands. Symmetric padding is applied to ensure that the embedded sequence maintains the same length as the input, thereby preventing positional offsets. In addition, the bias term is removed, while feature distribution normalization is achieved through a subsequent batch normalization layer, reducing parameter redundancy.

The primary role of the convolutional embedding layer is to transform the one-dimensional waveform signal into high-dimensional feature representations. This higher-dimensional space more effectively captures subtle differences in nucleotide hydrophobicity (e.g., the hydrophobicity difference of 0.50 between G and C), thereby providing a richer semantic foundation for subsequent positional encoding and multi-scale feature extraction. This design differs from the discrete embedding strategies commonly used in natural language processing while remaining well suited to the continuous nature of genomic signals. Moreover, the 1D convolutional embedding requires substantially fewer parameters and avoids large-scale lookup operations; its parameter count is only 0.03% of that of a conventional embedding layer, significantly reducing gradient computation and memory consumption during training.

Attention-enhanced positional encoding (WaveEncoder). Genomic signal function is strongly associated with positional context—for example, the ~50 bp region surrounding the transcription start site (TSS) constitutes a core regulatory zone, while the functional influence of distal bases decays substantially. Conventional fixed positional encoding schemes (e.g., linear bias or sinusoidal encoding) cannot dynamically adapt positional weights to different functional regions. To address this limitation, the model introduces the WaveEncoder module, which implements attention-enhanced positional encoding. Built upon the high-dimensional features generated by the convolutional embedding layer, this module dynamically integrates positional semantics with signal features.

The module first generates sinusoidal positional encodings with dimensions matching the feature space, parameterized by learnable frequency and phase variables. The periodic nature of the sinusoidal function aligns with the decay pattern of positional influence in genomic sequences, while the learnable parameters allow the encoding to adapt to positional characteristics across different species and functional regions (e.g., differences in the positional influence range of human versus yeast promoters). The module then concatenates the high-dimensional signal features with the positional encodings and learns attention weights through two successive 1D convolutional layers (kernel size: 15 bp, covering typical regulatory motifs and their adjacent regions). A Sigmoid activation function is subsequently applied to constrain the weights within the range [0,1]. The attention weights dynamically determine the contribution of signal features and positional encodings. In key signal regions (e.g., promoter core regions containing a TATA box), the weights favor signal features to prioritize biologically relevant information. In non-critical regions (e.g., distal non-regulatory segments), the weights favor positional encoding, allowing positional semantics to supplement redundant signal information. This dynamic fusion mechanism avoids the rigidity of fixed positional encodings while ensuring deep coupling between positional information and signal features.

Collaborative Optimization of InceptionTime and Dilated Convolutions. Genomic functional features span a wide range of scales—from transcription factor binding sites as short as 3 bp to enhancer core regions extending up to 60 bp. Single-scale convolutional extraction is therefore insufficient to capture the full spectrum of functional information, while simply enlarging convolutional kernel sizes can lead to parameter explosion and reduced computational efficiency. After the attention-based positional fusion stage, the model incorporates an InceptionTime–dilated convolution module. Through a design consisting of bottleneck compression, multi-branch feature extraction, and residual connections, the module enables efficient capture of multi-scale functional features.

The module first employs a 1×1 convolution to compress the 128-dimensional features into a 32-dimensional bottleneck representation, substantially reducing the computational complexity of subsequent branches. Batch normalization and the GELU activation function are applied to optimize feature distribution and enhance the model's nonlinear representational capacity. Subsequently, three parallel branches are introduced to achieve multi-scale coverage. The short-scale branch uses a 3 bp convolutional kernel (dilation rate = 1), focusing on local features such as single-nucleotide variations and short motifs. The long-scale branch employs a 21 bp convolutional kernel (dilation rate = 3), which expands the receptive field to 61 bp through dilation, enabling coverage of long-range regulatory regions without increasing kernel size. A global pooling branch further complements the representation by applying adaptive average pooling to capture global background characteristics (e.g., GC content distribution trends), preventing local feature extraction from overlooking broader contextual patterns. The outputs of the three branches are concatenated along the channel dimension and then compressed to 64 dimensions via a 1×1 convolution. A residual connection is introduced by projecting the pre-bottleneck 128-dimensional features to 64 dimensions and adding them to the branch outputs, thereby mitigating gradient vanishing in deeper networks. This design achieves non-redundant coverage across a scale range of 3–61 bp. Although it increases the parameter count by only about 15% compared with the conventional InceptionTime architecture, it simultaneously captures both local genomic details and global sequence trends.

Deep Integration of InceptionTime and Bidirectional GRU. The features extracted by the InceptionTime–dilated convolution module are fundamentally spatial multi-scale representations, where the channel dimension encodes information from different receptive scales. However, genomic function is determined by the sequential arrangement of nucleotides—for example, the ordering of specific motifs governs transcription factor binding affinity. Moreover, the reverse-complement nature of double-stranded DNA requires the model to capture bidirectional temporal dependencies along both the 5′→3′ and 3′→5′ directions. To address this requirement, the model introduces a fusion mechanism that integrates InceptionTime with a bidirectional GRU, transforming spatial multi-scale features into temporally structured representations.

First, the 64-dimensional spatial features produced by InceptionTime (with dimensions batch size × 64 × sequence length) are transposed into a sequence-first format (batch size × sequence length × 64) suitable for the bidirectional GRU, ensuring that multi-scale information is fully propagated into the temporal modeling module. The bidirectional GRU adopts a two-layer architecture: the first layer focuses on local temporal dependencies (e.g., interactions between adjacent motifs), while the second layer captures long-range temporal relationships, such as cooperative interactions among regulatory elements across different scales. Meanwhile, the forward GRU models dependencies along the 5′→3′ direction, while the backward GRU captures patterns along the 3′→5′ direction. Their outputs are concatenated along the hidden dimension (64 × 2 = 128), enabling comprehensive modeling of bidirectional sequence dependencies in double-stranded DNA. To further refine the temporal feature representation, a fusion layer processes the terminal GRU output (which encodes global sequence-level temporal information). This layer compresses the 128-dimensional feature vector to 64 dimensions using a two-layer fully connected network, with batch normalization, GELU activation, and dropout (rate 0.3–0.4) applied between layers. This fusion mechanism allows the spatial multi-scale advantages of InceptionTime to complement the temporal modeling capability of the GRU, effectively overcoming the limitations of relying solely on spatial or temporal modeling.

**Wavelet decomposition:**

**Level 1 decomposition:**

Approximation coefficients: $cA_1[k] = \sum_{n=0}^{5900} x_{enhanced}[n]h[n-2k],\ k = 0,1,\dots,2999$

Detail coefficients: $cD_1[k] = \sum_{n=0}^{5900} x_{enhanced}[n]g[n-2k],\ k = 0,1,\dots,2999$

**Level 2 decomposition:** using $cA_1$, as the input signal

Approximation coefficients: $cA_2[k] = \sum_{n=0}^{2999} x_{enhanced}[n]h[n-2k],\ k = 0,1,\dots,1499$

Detail coefficients: $cD_2[k] = \sum_{n=0}^{2999} x_{enhanced}[n]g[n-2k],\ k = 0,1,\dots,1499$

**Level 3 decomposition:** using $cA_2$, as the input signal

**…**

### Supplementary Notes:

**File Format:** Standard HDF5, supporting hierarchical groups, compression, and metadata.

**File Structure:**

```
ecoli_waves.h5
├── train/                  # Training set
│   ├── class_0/data        # Class 0 samples (float32 array, shape=[num_samples, 2000])
│   ├── class_1/data
│   └── ...
├── val/                    # Validation set
│   ├── class_0/data
│   └── ...
├── test/                   # Test set
│   ├── class_0/data
│   └── ...
└── Attributes
    ├── creation_date       # File creation date
    ├── author              # Author
    ├── description         # File description
    ├── dimensions          # Length of a single sample
    ├── split_ratio         # Train/val/test split ratio
    ├── class_labels        # Class names, corresponding to input FASTA files
    └── wavelet_params      # Wavelet compression parameters (wavelet/bank, decomposition
level, high-frequency threshold)
```

### Supplementary Figures:

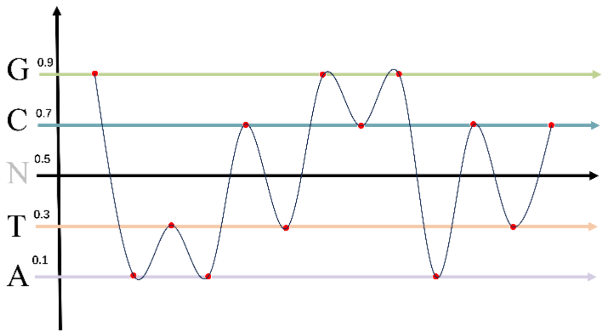


**Fig.S1. Schematic Illustration of Numerical Mapping.**

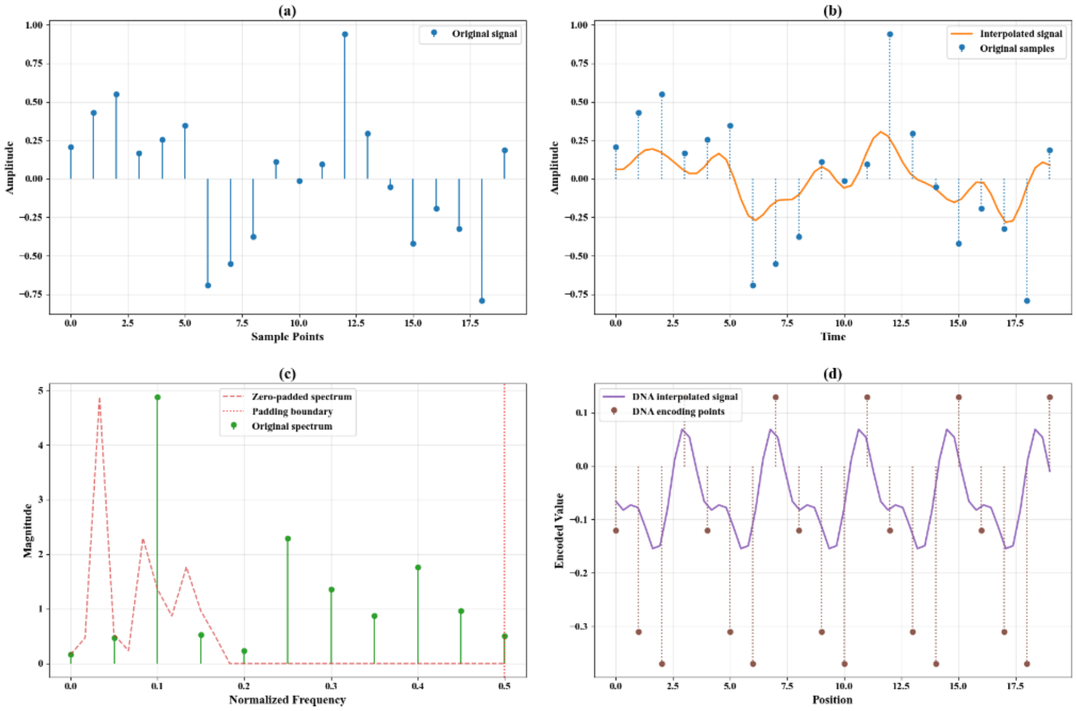


**Fig.S2. Schematic illustration of the Fourier frequency-domain zero-padding interpolation process.** (a) Original discrete hydrophobicity-encoded signal; (b) continuous waveform obtained through Fourier interpolation; (c) zero-padding operation in the frequency domain, with the red dashed line indicating the zero-padding boundary; (d) example of the continuous representation of the hydrophobicity index for a DNA sequence.

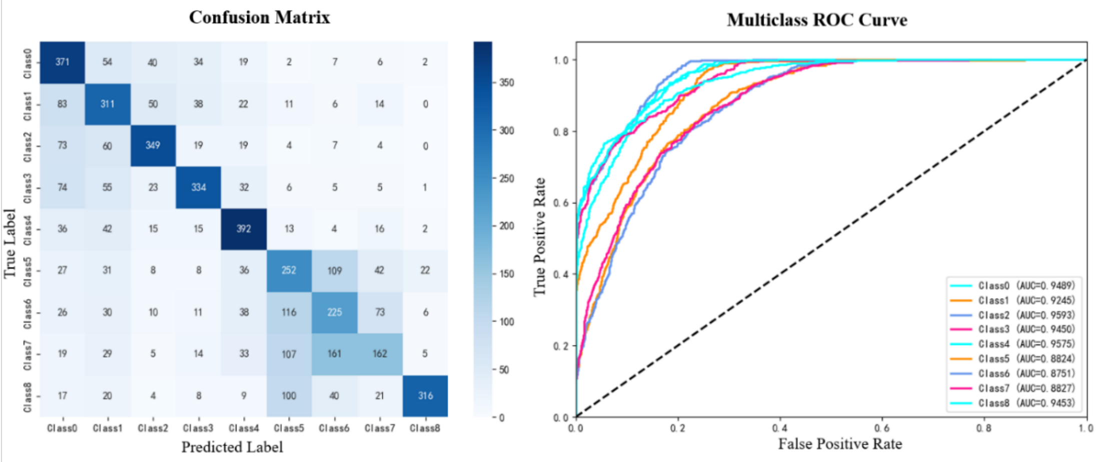


**Fig.S3. Confusion Matrix and ROC Curve for SARS-CoV-2 Mutation Classification**

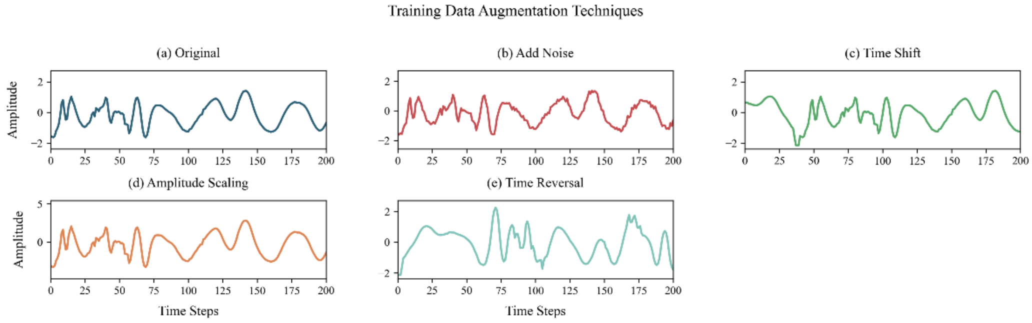


**Fig.S4. Illustration of biological signal augmentation strategies during the training phase.** (a) Original signal; (b)–(e) Robustness is enhanced via additive noise (SNR reduced by 8 dB), temporal shifting (±60 steps), amplitude scaling (0.8 –1.2×), and signal inversion (applied with 30% probability). More than 92% of key region features are preserved across all augmentations.

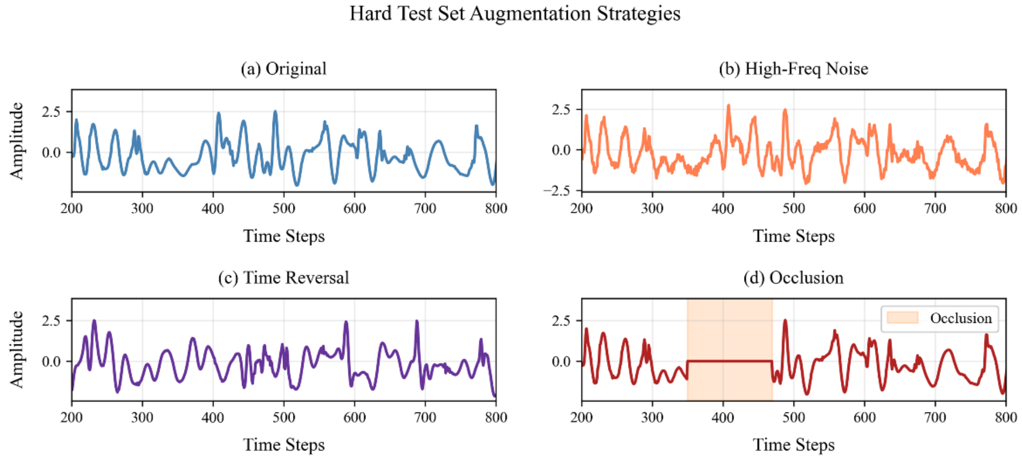


**Fig.S5. Simulation of signal perturbations used in challenge test augmentation.** (a) Original signal; (b)–(d) Challenge test augmentations introduce high-frequency noise (15% total signal energy), forced inversion, and random masking (50–150 steps) to simulate real-world perturbations such as sensor malfunction or data loss. These augmentations are used to evaluate the model's generalization under adverse conditions.

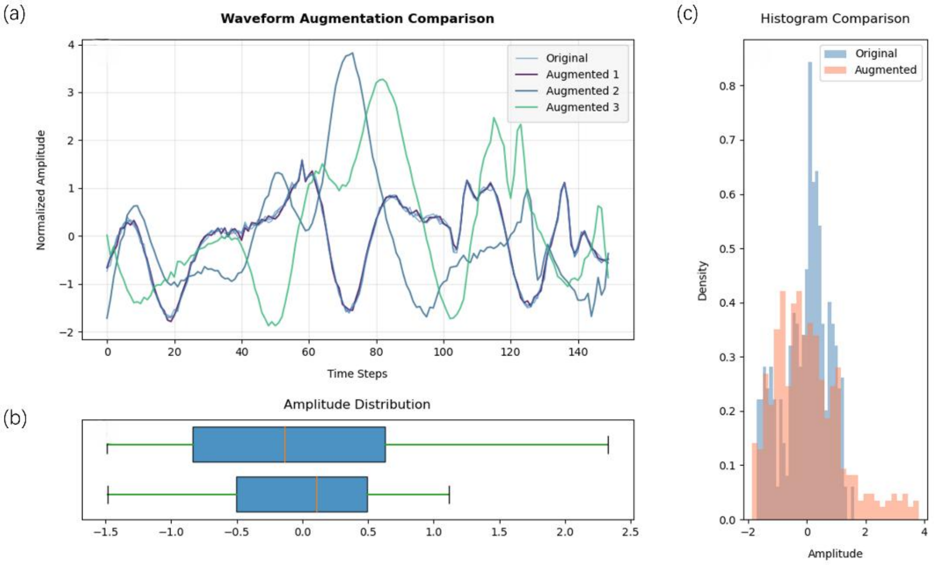


**Fig.S6. Multidimensional evaluation of hybrid data augmentation effects.** (a) The main waveform illustrates augmented signals (purple/green/cyan) with added noise, ±30% amplitude perturbations, and time shifts within 0–150 time steps. The shape integrity of key peaks is preserved with >95% fidelity. (b) Boxplot reveals a 38% increase in amplitude interquartile range (IQR) after augmentation, indicating an effective broadening of the data distribution. (c) The histogram shows that the augmented data (orange) increases probability density by 47% in the range [–2, 4], surpassing the original distribution (blue), which is constrained to [–2, 1.5]. This demonstrates that controlled perturbations enhance model robustness while preserving the temporal characteristics of the biological signals.

### Supplementary Tables:

Table S1.Evaluation metrics used in this study.

| Metric | Formula |
|---|---|
| Balanced Accuracy(BA) | $Balanced\ Accuracy = \frac{1}{C}\sum_{i=1}^{C}\frac{TP_i + TN_i}{TP_i + TN_i + FP_i + FN_i}$ |
| Macro-F1 | $Macro - F1 = \frac{1}{C}\sum_{i=1}^{C} F1_i$ <br> $F1_i = 2 \times \frac{Precision_i \times Recall_i}{Precision_i + Recall_i}$ <br> $Precision_i = \frac{TP_i}{TP_i + FP_i}$ <br> $Recall_i = \frac{TP_i}{TP_i + FN_i}$ |
| AUC-ROC | $\begin{cases} \text{Binary classification: AUC } = \text{ area under the ROC curve.} \\ \text{Multiclass classification: Macro} - \text{AUC} = \frac{1}{C}\sum_{i=1}^{C} AUC_i \end{cases}$ <br> where the horizontal axis of the ROC curve is $FPR_i = \frac{FP_i}{FP_i+TN_i}$ and the vertical axis is$TPR_i = \frac{TP_i}{TP_i+FN_i}$ |
| 95% CI | $95\%CI = [Precentile_{2.5}(\hat{\theta}_b), Precentile_{97.5}(\hat{\theta}_b)]$ |

Note: $C$ denotes the total number of classes in the task. $TP_i$, $TN_i$, $FP_i$, and $FN_i$ represent the true positives, true negatives, false positives, and false negatives, respectively, when the $i-th$ class is treated as the positive class (the subscript $i$ is omitted for binary classification tasks). $\hat{\theta}_b$ denotes the balanced accuracy estimate obtained from $n_{bootstrap} = 1000$ bootstrap resamples, and $Precentile_k(.)$ indicates the $k-th$ percentile. All metrics were implemented using the Python scikit-learn library to ensure reproducibility of the results.

Table S2. Experimental validation of numerical mapping.

| Nucleotide Order | Result | |
|---|---|---|
| | ACC | AUC-ROC |
| ATCG, GCTA | **0.6597 (95% CI: 0.6501~0.6700)** | **0.7195** |
| ATGC, CGTA | **0.6576 (95% CI: 0.6477~0.6669)** | **0.7107** |
| ACTG, GTCA | 0.6144 (95% CI: 0.6044~0.6247) | 0.6506 |
| ACGT, TGCA | 0.6230 (95% CI: 0.6131~0.6339) | 0.6660 |
| AGTC, CTGA | 0.6328 (95% CI: 0.6226~0.6425) | 0.6813 |
| AGCT, TCGA | 0.6404 (95% CI: 0.6310~0.6510) | 0.6838 |
| TACG, GCAT | **0.6626 (95% CI: 0.6530~0.6721)** | **0.7164** |
| TAGC, CGAT | **0.6522 (95% CI: 0.6423~0.6617)** | **0.7050** |
| TCAG, GACT | 0.6166 (95% CI: 0.6062~0.6264) | 0.6662 |
| TGAC, CAGT | 0.6042 (95% CI: 0.5939~0.6144) | 0.6601 |
| CATG, GTAC | 0.6362 (95% CI: 0.6261~0.6464) | 0.6845 |
| CTAG, GATC | 0.6431 (95% CI: 0.6334~0.6520) | 0.7032 |

Note: GC and AT nucleotides inherently possess distinct chemical structures, and GC content is closely associated with functional regions and fragmentation patterns in many genomic analysis tasks. The above experiments indicate that the model achieves higher performance when separating the AT group from the GC group, suggesting that the model partially leverages the "GC preference" sequence statistical feature for classification. When the numerical encoding captures the intrinsic physicochemical properties of nucleotides, the model can more effectively learn the mapping between sequences and their functional attributes.

Table S3. Data augmentation parameters across training and testing stages and their biological relevance analysis.

| Enhancement Stage | Enhancement Type | Technical Parameters | Biological Significance |
|---|---|---|---|
| Training Set Routine | Random Amplitude Scaling | Scaling Range: 0.8-1.2 (uniform)<br>Probability: 60% | Simulate signal intensity changes at different sequencing depths. |
| | Gaussian Noise Injection | Standard Deviation: 0.05 (SNR=26dB)<br>Probability: 80% | Simulate random errors in PCR amplification. |
| | Random Time Shift | Maximum Shift: ±60 bp<br>Probability: 40% | Simulate the uncertainty of read alignment and localization. |
| | Time Series Reversal | Reversal Probability: 30% | Simulate complementary strand sequencing scenarios. |
| Challenge Test | High-Frequency Noise | Standard Deviation: 0.10 (SNR=20dB) | Simulate sequencing noise in severely degraded samples. |
| | Random Time Reversal | Reversal Probability: 40% | Test the robustness of the model to sequence directionality. |
| | Random Local Occlusion | Occlusion Length: 100 - 300 bp<br>Position: 200 - 1600 bp<br>Probability: 60% | Simulate low-coverage regions in sequencing. |

Note: Some augmentation parameters are dynamically adjusted based on sequence length.
